# Evolutionary formation of a human *de novo* open reading frame from a non-primate non-coding genomic region that proceeds via biased random mutations

**DOI:** 10.1101/2023.08.09.552707

**Authors:** Nicholas Delihas

## Abstract

Two open reading frames (ORFs) of evolutionary interest stem from the human gene *SMIM45*. An investigation shows that one consists of an ultra-conserved 68 amino acid (aa) sequence that dates back to the amphibians, ∼192 MYA. In contrast, the other, an ORF of 107 aa develops slowly during primate evolution. An early developmental stage of the 107 aa ORF has been detected in non-primate genomes. In the mouse, it consists of a small sequence homologous to a segment of the human 107 aa ORF, the aa sequence SGLE**\***VTVYGGGVQKGKT. This suggests the evolutionary root of the 107 aa ORF dates to an ancestor of the mouse; root species have been proposed based on the presence or absence of the early developmental sequence in ancestors. The formation of the complete 107 aa ORF during primate evolution is also outlined. Found is growth of the 107 aa ORF by biased random mutations. The delineation of mutations occurring during development of the 107 aa ORF can provide a model for the evolutionary formation of ORFs of *de novo* protein genes.

## Introduction

The human gene *SMIM45* is unusual and complex. It encodes an evolutionary ultra-conserved 68 amino acid (aa) ORF (The UniProt Consortium 2023). This ORF is found next to an 107 aa ORF, a sequence that is not present in ancestors (Sayers 2022). Computational analysis shows that an *SMIM45* RNA transcript carries both the 68 aa and 107 aa ORF nt sequences. The transcript is transcribed and there is evidence for translation of the 68 aa ORF based on Ribo-Seq ribosome profiling, however the 107 aa sequence is not translated (NML/NCBI (Sayers 2022) (see Methods for online source). No function for the 68 aa protein has been presented but in an interesting study, An and coworkers (2023) describe a protein that functions in human brain development and predict that it is the 107 aa ORF of *SMIM45*. Aside from the question of function, the two contrasting ORFs offer an interesting study in the divergent evolutionary history of *SMIM45*.

The work presented here traces the evolutionary pattern of development of the *SMIM45* ORFs with a primarily analysis of the 107 aa ORF. The NIH Mammalian Gene Collection Program that identifies cDNA clones that contain complete ORFs for human and mouse genes (Strausberg et al. 2002) was crucial to this study. Described is use of genomic DNA sequences and their translated ORFs to analyze the evolutionary development of the *SMIM45* locus. The genomic DNA provides important information on ORF development and how particular regions evolve over evolutionary time. A protocol is described that is used to analyze development of the 107 aa ORF. The protocol can be a model for the analysis of ORF development from human *de novo* protein genes, as one would expect that the creation of a *de novo* gene sequence begins with formation of its ORF. *De novo* protein genes are genes that originate from a non-coding genomic region of an ancestral genome, i.e., “by scratch”. Contrary to past views that new genes are primarily formed by gene duplication (Ohno 1969), a large body of evidence accumulated during the past 15-20 years shows that a significant number of protein genes originate by *de novo* means (Long M. et al. 2003; Levine et al. 2006; Zhou and Wang 2008; Carvunis et al. 2012; Wu and Zhang 2013; Tautz D. 2014; Schlötterer C. 2015; Schmitz and Bornberg-Bauer 2017; Van Oss et al. 2019.; Vakirlis et al. 2020; Broeils et al. 2023).

Non-coding regions that generate *de novo* genes can be visualized as ancestral DNA that contains random sequences, but with an initial start of a protein ORF that involves a selected root base pair (bp) or bps. The detection of the evolutionary root of an ORF is exceedingly difficult, as distinguishing root DNA base-pairs from random sequences is very problematic and attempts to find root bps in ancestral genomic DNA that has no coding capacity have not been reported. However, the evolutionary progression of an ORF that carries a transmembrane motif from an ancestral non-coding sequence has been described for the yeast genomic locus, YBR196C-A (Vakirlis et al. 2020), and Vakirlis et al. 2022) reported the evolutionary formation of a class of microproteins in primates where early developmental stages and species of origin have been proposed.

In the work shown here on the evolutionary development of the human *SMIM45* 107 aa ORF, found is a small nt sequence in the mouse genome that is homologous to a segment of the human 107 aa mRNA and which translates to the aa sequence SGLE**\***VTVYGGGVQKGKT; this represents an early developmental stage of the107 aa sequence. However, ancestral species were also analyzed for possible root origins of the 107 aa sequence, and proposed is initial formation of the early developmental sequence in a species that evolutionarily appeared before the elephant (103 million years ago (MYA) but after the koala (164 MYA).

With the use of theoretical modeling, Thibault and Landry (2019) proposed that biased mutations are a driving force in birth of *de novo* genes. Mathematical modeling has also been used to investigate the formation of *de novo* ORFs by biased mutations and to address the question of how quickly genes emerge and are lost (Lyengar and Bornberg-Bauer 2023). Reported here is the mutational process that occurs during the evolution of the 107 aa ORF in primates. Found is the formation of the ORF by biased random mutations, i.e., the biased selection of a bp from a mutation where the bp is fixed during evolution but with random mutations also occurring during the evolutionary process. Presented here is an analogy of the biased random mutational process to a biased random walk that occurs during chemotaxis (Macnab and Koshland K Jr 1972; Sourjik V, and Wingreen 2012).

## Methods and Materials

### Protocol for finding the evolutionary early developmental stage of the 107 aa ORF

1. The genomic region of the mouse *SMIM45* that contains the 68 aa ORF displays synteny with the genomic region containing the human 107 aa ORF nt sequence. This serves as a guidepost for locating genomic nt sequences that are homologous to the human 107 aa nt sequence in the mouse and other species. By aligning the *SMIM45* gene sequence from the mouse and other species with the sequences of human *SMIM*45 exon 2 and the human 107 aa mRNA, the mouse *SMIM45* region that has homology with the human 107 aa mRNA is determined. The EMBL-EBI Clustal Omega alignment program https://www.ebi.ac.uk/Tools/msa/clustalo/) is used for nt and aa sequence alignments. When studying a human gene that is not annotated in an ancestral species, the entire genomic sequence between two flanking genes that display synteny can be analyzed for similarities with the gene nt sequence. This is described below for *SMIM45*. Important are adjacent evolutionary conserved sequences to use as guideposts. With other genes, highly conserved enhancer sequences, when found adjacent to or within the gene being analyzed can also be useful as a guidepost in locating a homologous sequence in an ancestor. With *SMIM45*, *SEPTIN3* served as the consistent guidepost.
2. A segment of the mouse *SMIM45* nt sequence that produces a stretch of similarity to the human 107 aa mRNA is used to translate the nt sequence by the Expasy translation tool. One of the resulting aa sequences from the three translated 5’3’ Frames is found to contain the early developmental sequence. Thus, by aligning the translated three 5’3’ Frame sequences from the mouse or primate species with the human 107 aa ORF aa sequence, the early developmental aa sequence homologous to the human 107 aa sequence is obtained.
3. For primates, as the growing aa sequence evolves during evolution of the ORF, the 5’3’ Frame aa alignments can identify additional aa that have an identity with the 107 aa sequence. With the mouse genomic sequence, only random identities are found outside of the early developmental aa sequence.
4. For species that do not have the *SMIM45* gene annotated (e.g., the tarsier), the *SMIM45* sequence was determined by analyzing the DNA sequence between the two genes that show synteny, *SEPTIN3* and *CENPM*. Nt sequences of the genomic regions are then aligned with the 68 aa sequence. This gave an accurate location of the homologous 107 aa nt sequence and/or the early embryonic sequence in these species.
5. To assess possible species that may represent root origins of the 107 aa ORF, entire *SMIM45* gene sequences from various species were aligned with the nt sequences of the 68 aa, 107 aa, and the early developmental sequence SGLELVRVCGGGMQRDKT. Genomes of ancestral species that show no identity to the early developmental sequence, and species representing the closest related progeny and which display a similarity to the early developmental sequence as well as showing synteny with the 68 aa sequence were guides in predicting root origins.

### Species analyzed for 68 aa and/or 107 aa ORFs

*Bombina bombina*, (the fire-bellied toad), *Gallus gallus* (chicken), *Ornithorhynchus anatinus* (platypus), *Phascolarctos cinereus* (koala), *Monodelphis domestica*, (gray short-tailed opossum), *Elephantulus edwardii* (Cape elephant shrew), *Elephas maximus indicus* (elephant), *Talpa occidentalis* (Iberian mole), *Bos taurus* (cattle), *Leopardus geoffroyi* (Geoffroy’s cat). *Oryctolagus cuniculus* (rabbit), *Mus musculus* (house mouse), *Tupaia chinensis* (Chinese tree shrew), *Lemur catta* (Ring-tailed lemur), *Carlito syrichta* (Philippine tarsier), *Macaca mulatta* (Rhesus monkey), *Macaca thibetana thibetana (*The Tibetan macaque)*, Papio anubis* (olive baboon), *Pongo abelii* (Sumatran orangutan), *Gorilla gorilla gorilla* (western lowland gorilla), *Pan troglodytes* (chimpanzee), and *Homo sapiens* (human).

### Source and properties of gene and transcript sequences

The *SMIM45* gene sequence was obtained from the website: home gene NCBI, https://www.ncbi.nlm.nih.gov/gene/?term=smim45+human (Sayers et al. 2022. The exon 2 sequence is from *Ensembl* https://useast.ensembl.org/Homo_sapiens/Gene/Sequence?db=core;g=ENSG00000205704;r=22:41952150-41958939;t=ENST00000381348 (Cunningham et al. 2022.) The *SMIM45* protein data were from UniProtKB/Swiss-Prot (The UniProt Consortium 2023) and the NLM/NCBI (Sayers et al. 2022.).

The *SMIM45* exon 2 nt sequence is:

> 1 ccaaggccgc cgcgatgccg cacttcctgg actggttcgt gccggtctac ttggtcatct
>
> 61 cggtcctcat tctggtgggc ttcggcgcct gcatctacta cttcgagccg ggcctgcagg
>
> 121 aggcgcacaa gtggcgcatg cagcgccccc tggtggaccg cgacctccgc aagacgctaa
>
> 181 tggtgcgcga caacctggcc ttcggcggcc cggaggtctg agccgacttg caaaggggat
>
> 241 aggcgggcgg caccgggcgc cctcccccag cccgccccgc ccgcccagcc cggagacccc
>
> 301 caaggcagag ggaggccggc ctgttggccc tccacgctat ccctctgcag cctgggccct
>
> 361 cccgacagag gccccaggtg cgctggcagt ggaggtgggg cacttaggtg cctggctggc
>
> 421 ccagggcttg ctctccgtgt caagccgact cacccagagc ccaccctccc aagctcaggg
>
> 481 gcatcctccg ctgggcccca gtgcctttgc gctgcgcagc actctgccct ccactggact
>
> 541 caggcatgtc tatggctgcc tgtcctgagg ctccggagcc ctcatttctt cgtgaagtcc
>
> 601 ccagctcccc tgcctccact caatggcacc ggccctgcaa ctttaggcag gtcgaagcca
>
> 661 acccaaggaa agaacctaag aacctcgttt ggagggatgt cagcttgggc cagaccagcc
>
> 721 gcaccccgcg gggctcaggc ttggaactgg tgagggtgtg tggtgggggt atgcagaggg
>
> 781 ataagaccgt ggtagaggag agggttggtg aggagagaga gagagagaga gagagagtct
>
> 841 ggggggagcg ggcaagcatg gggagatgag atgtgtatat gtgagagaga gtgtgggggc
>
> 901 cccaggcagg gcaggaggtg gtggaaacgg ggtgaactcc gtgggctgtg tgaggactgt
>
> 961 ccatagtggg tcccaacccc ctccctctgc tggagtttcc tagcccttcc ccctccccaa
>
> 1021 gactgtggca gcaggcagga gcccctgccc tccctccctg tcctgtgcca cacttctggg
>
> 1081 gccaaaccca gcccccttga gccaggccct gccagactcc aagcccaccc tagaaccctc
>
> 1141 ctcctgtgtg gagactctgt tgccccactt tggacacaga ttggcaacct gcctcacccc
>
> 1201 gccccccttc gctggggctt ccatcttaat ttattctcaa taataaagac ttcatgatga
>
> 1261 tctctgca

The deduced mRNAs that encode the human 68 aa ORF and 107 aa ORF are:

68 aa mRNA

> 1 atgccgcact tcctggactg gttcgtgccg gtctacttgg tcatctcggt cctcattctg
>
> 61 gtgggcttcg gcgcctgcat ctactacttc gagccgggcc tgcaggaggc gcacaagtgg
>
> 121 cgcatgcagc gccccctggt ggaccgcgac ctccgcaaga cgctaatggt gcgcgacaac
>
> 181 ctggccttcg gcggcccgga ggtctga

107 aa mRNA

> 1 atgtctatgg ctgcctgtcc tgaggctccg gagccctcat ttcttcgtga agtccccagc
>
> 61 tcccctgcct ccactcaatg gcaccggccc tgcaacttta ggcaggtcga agccaaccca
>
> 121 aggaaagaac ctaagaacct cgtttggagg gatgtcagct tgggccagcc cagccgcacc
>
> 181 ccgcggggct caggcttgga actggtgagg gtgtgtggtg ggggtatgca gagggataag
>
> 241 accgtggtag aggagagggt tggtgaggag agagagagag agagagagag agtctggggg
>
> 301 gagcgggcaa gcatggggag atga

The 68 aa mRNA nt coordinates are positions 15-221 of *SMIM45* exon 2 and the 107 aa mRNA are positions 546-869. The 107 aa mRNA is a putative mRNA. The nomenclature used for the mRNAs is that shown by the NLM/NCBI.

The 68 aa ORF has a predicted alpha transmembrane domain (Supplemental File 1 Fig. S1) (Hallgren J. et al. 2022). Its base composition is:

A(15% 33) | T(22% 43) | G(31% 66) | C(32% 69)

The NML/NCBI computational analysis of *SMIM45* shows a transcript, NM_001395940.1, Homo sapiens small integral membrane protein 45 (SMIM45) transcript variant 1 mRNA. This transcript is identical to *Ensembl* transcript ENST00000381348.5 SMIM45-201. The transcript contains both the 68 aa and 107 aa mRNA sequences. UniProt provides evidence for transcription of ENST00000381348.5 (A0A590UK83 · SMI45_HUMAN). The NLM/NCBI and Ensembl list the 68 aa as a protein product based on the conservation of the ORF and Ribo-Seq elongating ribosome analysis, but there is no protein product for the 107 aa ORF ((https://www.ncbi.nlm.nih.gov/genome/gdv/browser/genome/?cfg=NCID_1_31487677_130.14.22.10_9146_1696530571_2728389868). It appears that there is no read-through of transcript NM_001395940.1 that would include the 107 aa ORF.

The nt sequence of transcript variant 1, mRNA NM_001395940.1 is:

> 1 gctctagggg ctggactcag ggcggtttga aagatcggcg cgcaccgcag gagcaacggt
>
> 61 tggtcctgcg gctgtgatgt cggtgttgag gcccctggac aagctgcccg gcctgaacac
>
> 121 ggccaccatc ttgctggtgg gcacggagga tgctcttctg cagcagctgg cggactcgat
>
> 181 gctcaaagag gactgcgcct ccgagctgaa ggtccacttg gcaaagtccc tccctttgcc
>
> 241 ctccagtgtg aatcggcccc gaattgacct gatcgtgttt gtggttaatc ttcacagcaa
>
> 301 atacagtctc cagaacacag aggagtccct gcgccatgtg gatgccagct tcttcttggg
>
> 361 gaaggtgtgt ttcctcgcca caggtgctgg gcgggagagc cactgcagca ttcaccggca
>
> 421 caccgtggtg aagctggccc acacctatca aagccccctg ctctactgtg acctggaggt
>
> 481 ggaaggcttt agggccacca tggcgcagcg cctggtgcgc gtgctgcaga tctgtgctgg
>
> 541 ccacgtgccc ggtgtctcag ctctgaacct gctgtccctg ctgagaagct ctgagggccc
>
> 601 ctccctggag gacctgtgag ggtggctggc ccctgggctg ccccttctca tggcttcgtg
>
> 661 ctgactccat aaacattctc tgttgaggat gtccagtcag ggcttgacag gcccaggctc
>
> 721 agcccgccgt ggctgggaag gttccctgca gtgccagtgc tgcagcaggg agagctgggc
>
> 781 agaagcagcg agggggccca gctggcgaga ctgtagcccc ctcccactcc cacactcact
>
> 841 cttgcagagc ctgtgtcttt aagcagctgg cgtgttacat ctccatttaa ggtttccttt
>
> 901 gaacaaaagg tctgtggcta aaaaaagttt aaaaatca

The theoretical RNA sequence that translates to the human conserved early developmental sequence (SGLELVRVCGGGMQRDKT) is:

tcaggcttggaactggtgagggtgtgtggtgggggtatgcagagggataagacc

The sequence is G rich: A(20%) T(23%) G(46%) C(11%)

### Evolutionary age of species

The evolutionary ages of primates and pre-primate species were from (Shao el al. (2023; Kuderna el al. 2023; Siepel, 2009). The evolutionary age of *Bombina bombina*, (the fire-bellied toad) was from (Feng et al. 2017), the cow from Zhang et al. (2010)., koala from (Phillips and Fruciano 2018).

### Nucleotide and amino acid sequence alignment and translation tools

Clustal Omega was employed to align three or more nt or aa sequences. Both EMBOSS Needle and Clustal Omega (Pairwise Sequence Alignment Tools<EMBL-EBI) were used for alignment of two sequences. In regions where the two sequences are similar, no significant differences were found between the two alignment programs. The data presented in Figures 4, and 6-8 show alignments by Clustal Omega. This was chosen over alignments by EMBOSS Needle as the Clustal Omega format readily allows for visual identification of aa positions of identity.

The MAFFT online service: multiple sequence alignment, interactive sequence choice and visualization (Katoh et al. 2019) was used to align early developmental sequences and visualize DNA base conservation with color coordination.

The ExPASy Translational tool (Gasteiger et al. 2002) was used for the translation of nucleotide sequences.

The Sequence Manipulation Suite (Stothard 2000) was used to obtain reverse complement sequences and for sequence clean up.

## Results

### Characterization of the human *SMIM45* gene and mouse *SMIM45* locus

In the work described here, it was possible to follow the evolution of DNA that leads to the formation of the human *SMIM45* 68 aa and the 107 aa ORFs because of the extreme evolutionary conservation of the 68 aa sequence. This sequence was used as a guidepost and in addition, the synteny displayed in the chromosomal region also served as a guidepost. *SMIM45* shows synteny with two neighbor genes, *CENPM* and *SEPTIN3*, with *SMIM45* situated in between (Fig. 1a, b).

**Fig. 1a, b.**
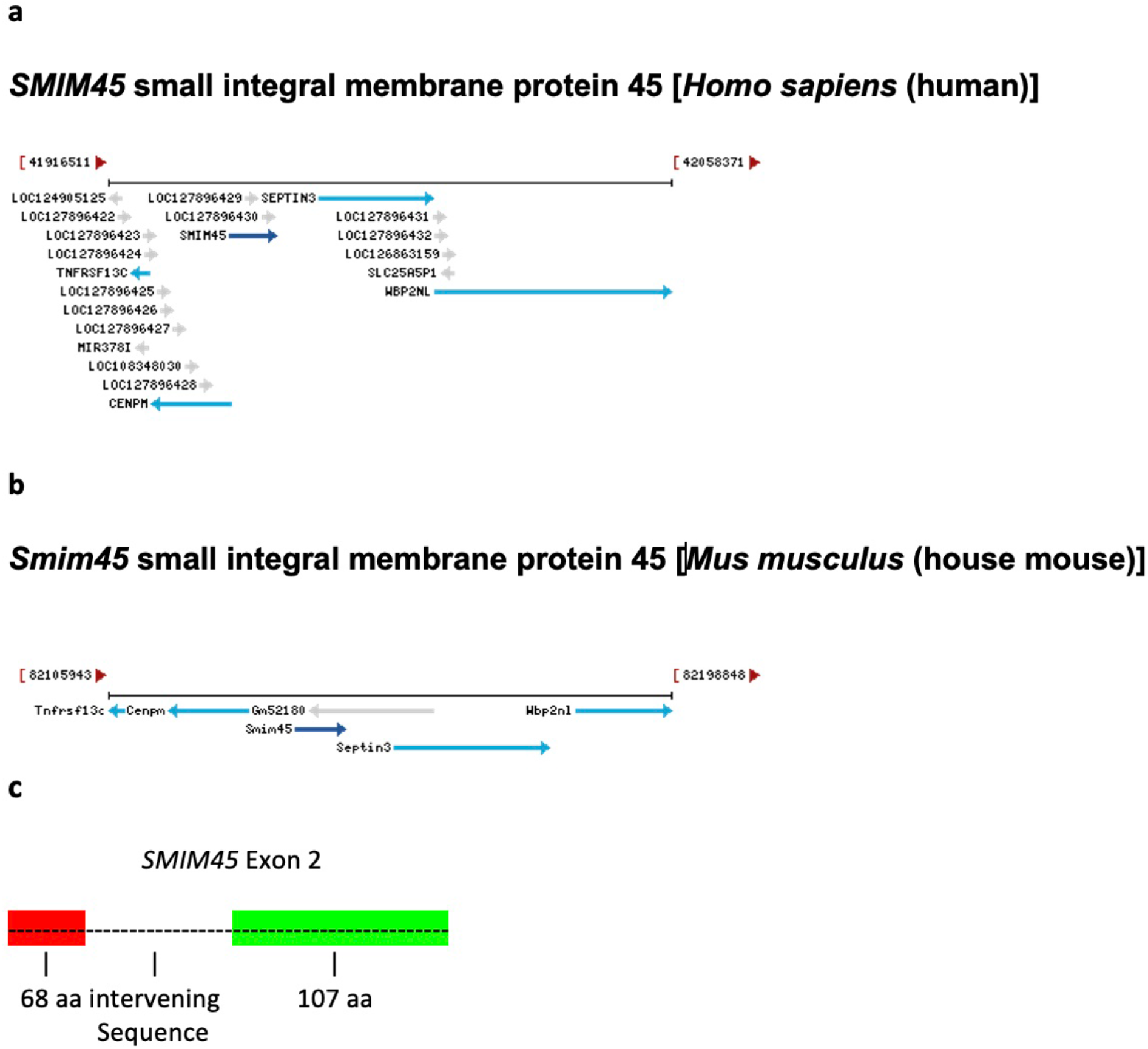
Chromosomal locations of *SMIM45* in the human and mouse genomes. The synteny with genes *CENPM* (centromere protein M) and *SEPTIN3* (neuronal-specific septin-3 gene) is shown. Diagrams are from the NLM/NCBI website that describes genes from different species (Home-Gene-NCBI, https://www.ncbi.nlm.nih.gov/gene). **1c.** A schematic of exon 2 of human *SMIM45* showing the arrangement of the 68 aa, the intervening sequence and the 107 aa sequence. The intervening sequence is 323 bp in length.

Two adjacent sequences are present in exon 2 of gene *SMIM45* and consist of two ORFs separated by a small intervening 323 bp sequence in human chromosome 22q11 (Fig. 1c); a translation of the intervening sequence shows that it is devoid of methionine and open reading frames.

To determine origins of *SMIM45* and how its sequence evolved, an analysis was performed on the evolutionary formation of the gene through its two ORFs, which constitute the translated sequences from the two segments of *SMIM45* exon 2 DNA. The 68 aa ORF displays a remarkable conservation over evolutionary time (Fig. 2). There is only one amino acid residue difference between the mouse (*Mus musculus*) and human sequence; the aa sequence from the tarsier [Philippine tarsier (*Carlito syrichta*)], a lower primate, shows 100% identity with the human sequence. The Philippine tarsier primate evolutionarily appeared ∼58 million years ago (MYA) and the mouse ∼87 MYA. This degree of ultra-conservation during primate evolution points to an important function of this ORF. However, we have also been able to trace the 68 aa sequence as far back as the amphibians (*Bombina bombina*, the fire-bellied toad), to an evolutionary age ∼192 MYA. Fig. 2 bottom shows an alignment of the toad sequence with that of the human 68aa and displays a 74% identity.

**Fig. 2.**
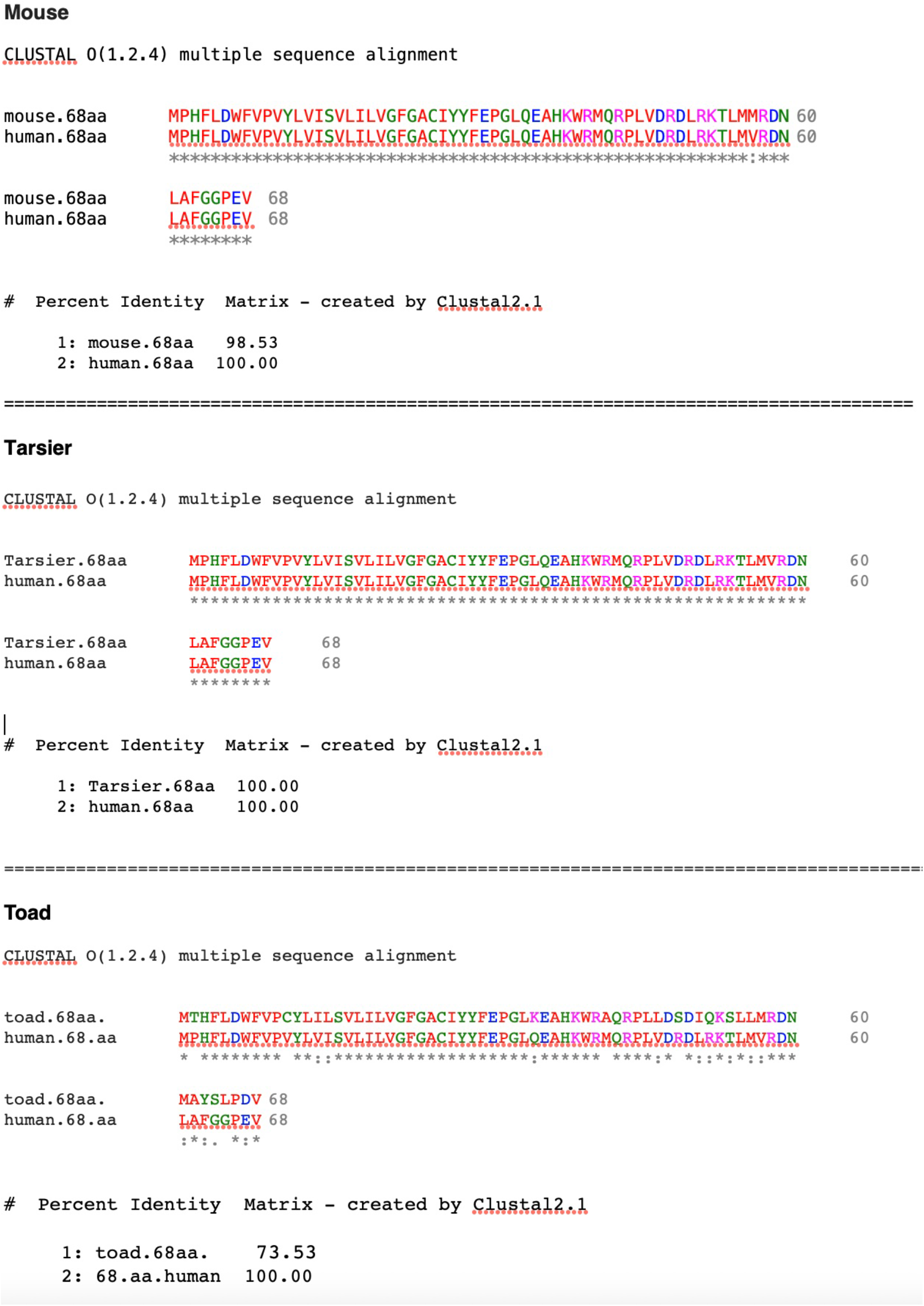
Alignment of the mouse 68aa ORF with the human 68 aa ORF (top), Philippine tarsier (*Carlito syrichta*) (middle) and fire-bellied toad (*Bombina bombina)* (bottom). The translated human and mouse *SMIM45* nt sequences are identical with the exception of one aa residue difference (position 57).

In sharp contrast to the 68 aa ORF, the 107 aa ORF evolved slowly over ∼100 million years of evolutionary time. The evidence presented suggests that a sequence representing an early developmental stage of the 107 aa sequence is in a mouse genomic region that has sequence similarity to the human107 aa nt sequence. An alignment of a segment of the mouse *SMIM45* gene sequence, a segment of exon 2, and the human mRNA that encodes the 107 aa ORF shows a 57.5% nucleotide sequence identity (Fig. 3), with one section, nt positions 7753-7810 of the mouse SMIM45 gene sequence displaying a 79.7% identity. The alignment locates the mouse genomic region that is homologous to the mRNA of the human 107aa.

**Fig. 3.**
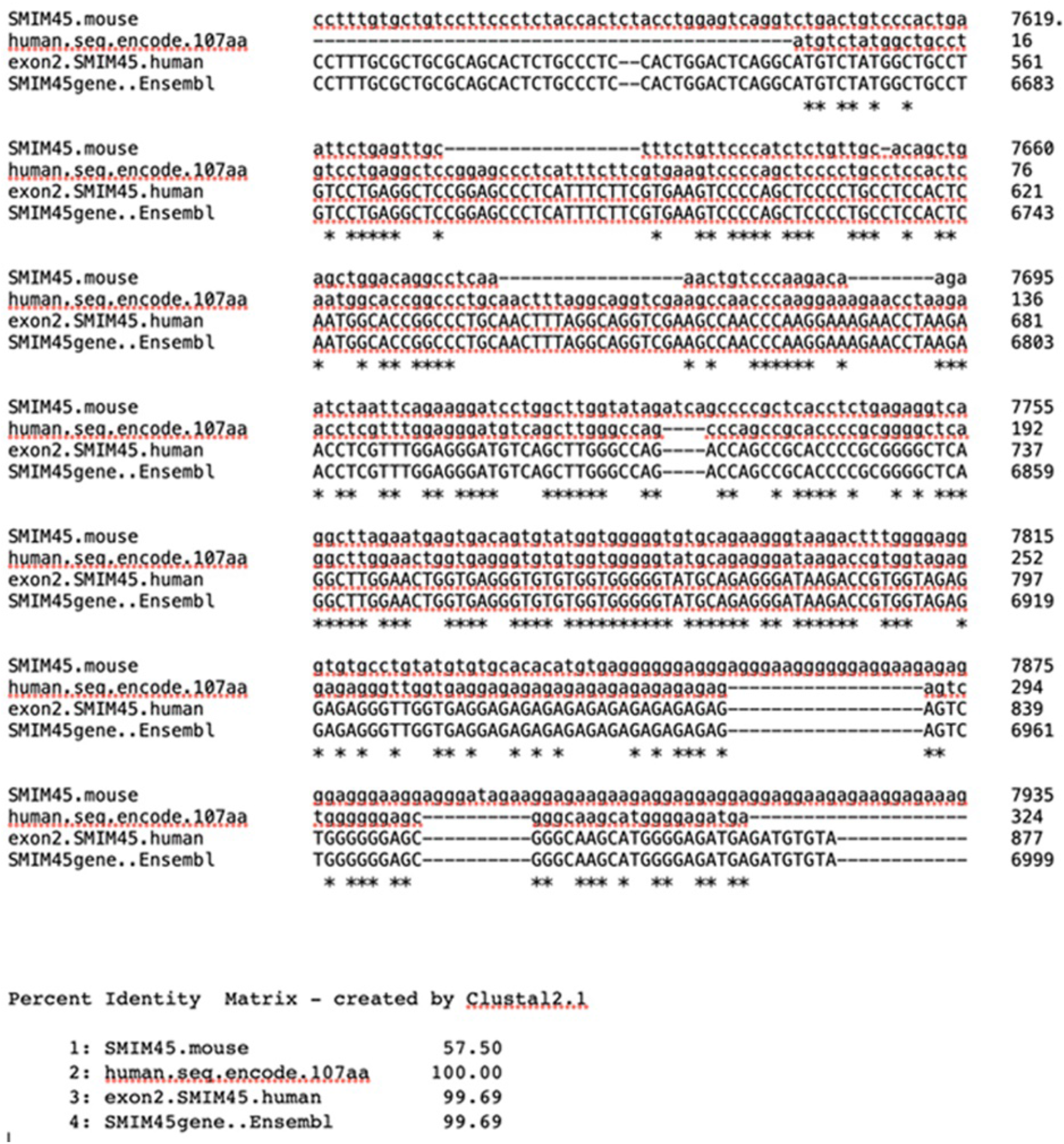
Alignment of a segment of the mouse *Smim45* gene sequence with the human sequence that codes for the 107 aa open reading frame, together with a segment of the nt sequence of exon 2 of the NLM/NCBI annotated human *SMIM45* gene and a segment of the human *SMIM45* gene sequence (as provided by *Ensembl*). The mouse *SMIM45* gene sequence is 8374 bp as annotated by the NCBI. The alignment was performed with the entire mouse *SMIM45* gene but only a segment of the sequence is shown in Figure 3.

A translation of the nucleotide (nt) sequence of human *SMIM45* exon 2 displays the 107 aa ORF, highlighted in light pink (107 aa) (Fig. 4a). However, a translation of the mouse genomic nt sequence homologous to the human 107 aa nt sequence encodes no ORFs with the exception of two methionine start codons followed by small aa sequences (highlighted in pink, Fig. 4b); these have no identity to the 107 aa ORF, and from Blast searches, do not form part of proteins in any organisms. A lack of a significant open reading frame indicates that the mouse homologous sequence is non-coding. However, found in the mouse genomic region is a nt sequence that translates to the aa sequence SGLE**\***VTVYGGGVQKGKT (**\*** represents a stop codon). The sequence is shown in Fig. 4b, highlighted in turquoise, 5’3’Frame 3. The homolog in humans is also highlighted in turquoise (Fig. 4a). The sequences flanking the turquoise highlighted sequence in the mouse genome appears to be random sequences as they do not show identity with proteins from any species.

**Fig. 4.**
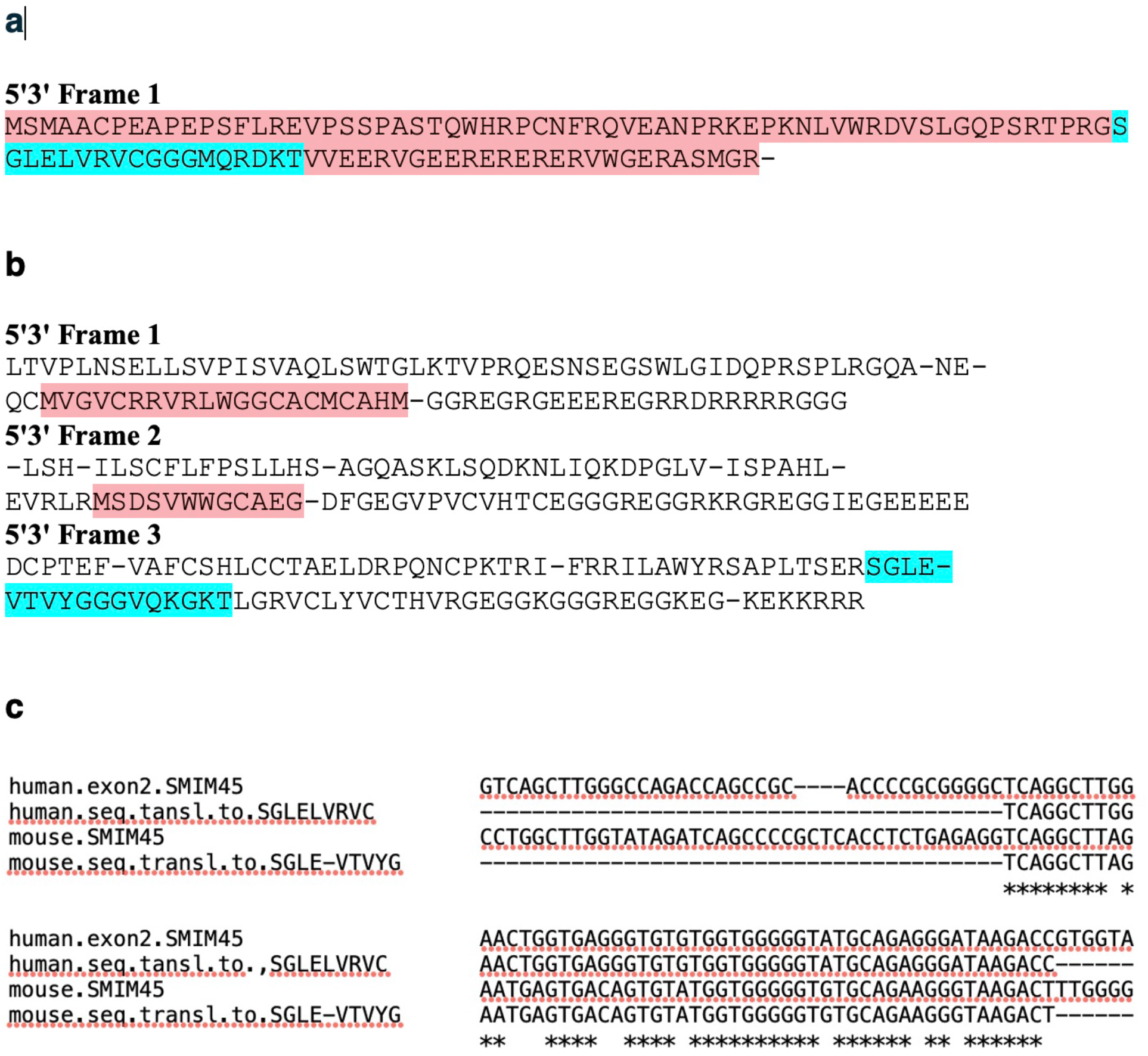
**a.** Translation of the nt sequence from human *SMIM45* exon 2 107aa ORF region. The 5’3’ Frame 1 is shown. The light pink highlighted sequence is the 107 aa open reading frame. **4b.** The translation of the mouse genomic region predicted to be homologous to the human 107 aa nucleotide sequence contains the sequence SGLE-VTVYGGGVQKGKT sequence (highlighted in turquoise). The highlighted sequences in light pink are small open reading frames (5’3’ Frames 1 and 3). **4c.** An alignment of nt sequences from the human exon 2 and mouse SMIM45 gene, together with the nt sequences that translate to the human aa sequence SGLELVRVCGGGMQRDKT and the mouse homologous aa sequence, SGLE**\***VTVYGGGVQKGKT (note in the alignment program that was used, the stop codons are shown with a dash; the star in the sequence shown here, conventionally represents a stop codon).

Fig. 4c shows an alignment of segments of nt sequences from the mouse *SMIM45* gene with that of the human *SMIM*45 exon 2, including nt sequences that translate to the human aa sequence SGLELVRVCGGGMQRDKT and the mouse homologous aa sequence, SGLE**\***VTVYGGGVQKGKT. The nt sequence that encodes the aa sequence segment in humans is 54 nt in length; there is an 81% identity between the human and the mouse homologous nt sequences. The probability of a random 54 nt sequence that has 81% identity by chance with the human counterpart, is exceedingly small, 1/(4^54^}xC(54,45)x3^9^. [C(54,45) is the binomial coefficient counting the ways to pick 45 positions out of 54 positions. The denominator is all sequences of 4 symbols of length 54, and 3^9^ represents ways to build sequences of non-matching nucleotides]. Thus, the mouse aa sequence, SGLE**\***VTVYGGGVQKGKT appears to represent a recognizable beginning of the formation of the human 107 aa open reading frame and an early stage of developmental in 107 aa open reading frame formation.

Fig. 5 is a graphic representation depicting the *mouse SMIM45* gene showing the genomic region comparable to the human 107 aa nt sequence, which includes the region encoding the early developmental aa sequence SGLE**\***VTVYGGGVQKGK flanked on both sides by random sequences.

**Fig. 5.**
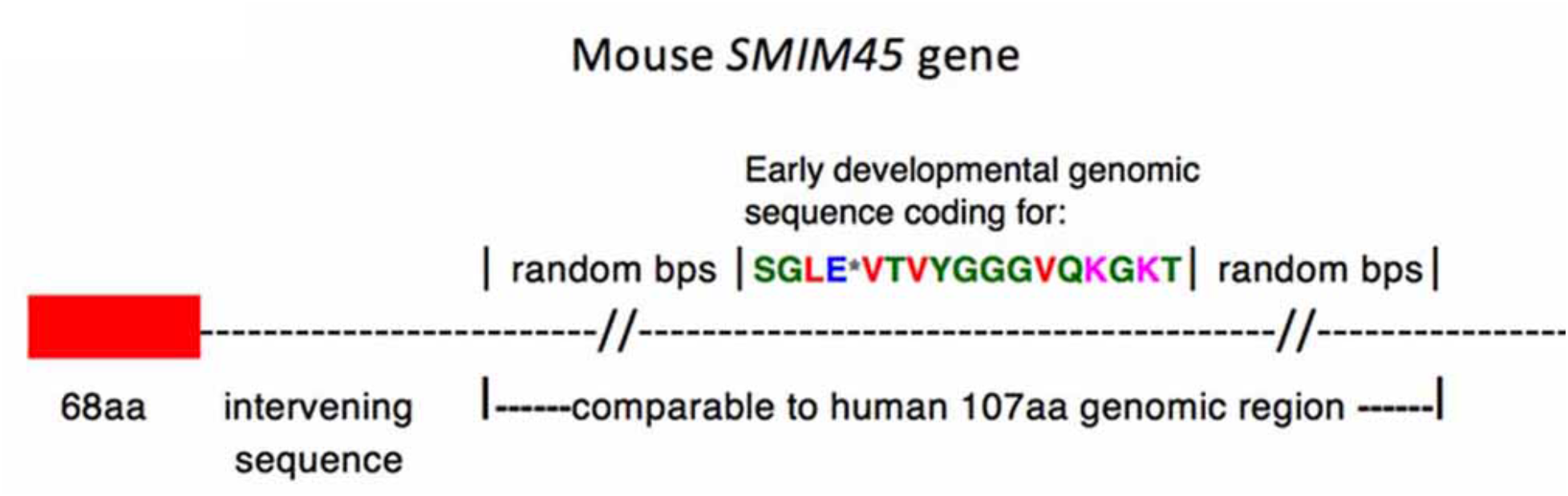
Graphic representation depicting the *mouse SMIM45* gene.

Table 1 shows the evolutionary progression of the early developmental sequence in primates. Development of the human sequence SGLELVRVCGGGMQRDKT proceeds rapidly; the lemur, a primitive and early evolutionary primate, formed an almost complete sequence corresponding to the human sequence, showing an 83% identity. There is a 100% similarity of the *Tibetan macaque* and the baboon (*Papio anubis)* sequences, both Old World monkeys, with the human sequence. However, the evolutionary pattern shows that sequences from several species deviate in similarity, the tree shrew, tarsier, and gorilla. The tree shrew sequence has a stop codon and several point mutations not present in the mouse sequence. These appear to be random mutations as they are not found in descendant species. The gorilla and tarsier sequences result from insertions/deletions that produce frame shift mutations. These mutations also appear to be of doubtful significance as they are not present in progenies. Thus, the formation of the human early development sequence was completed ∼31 MYA, but with random mutations occurring.

**Table 1.**
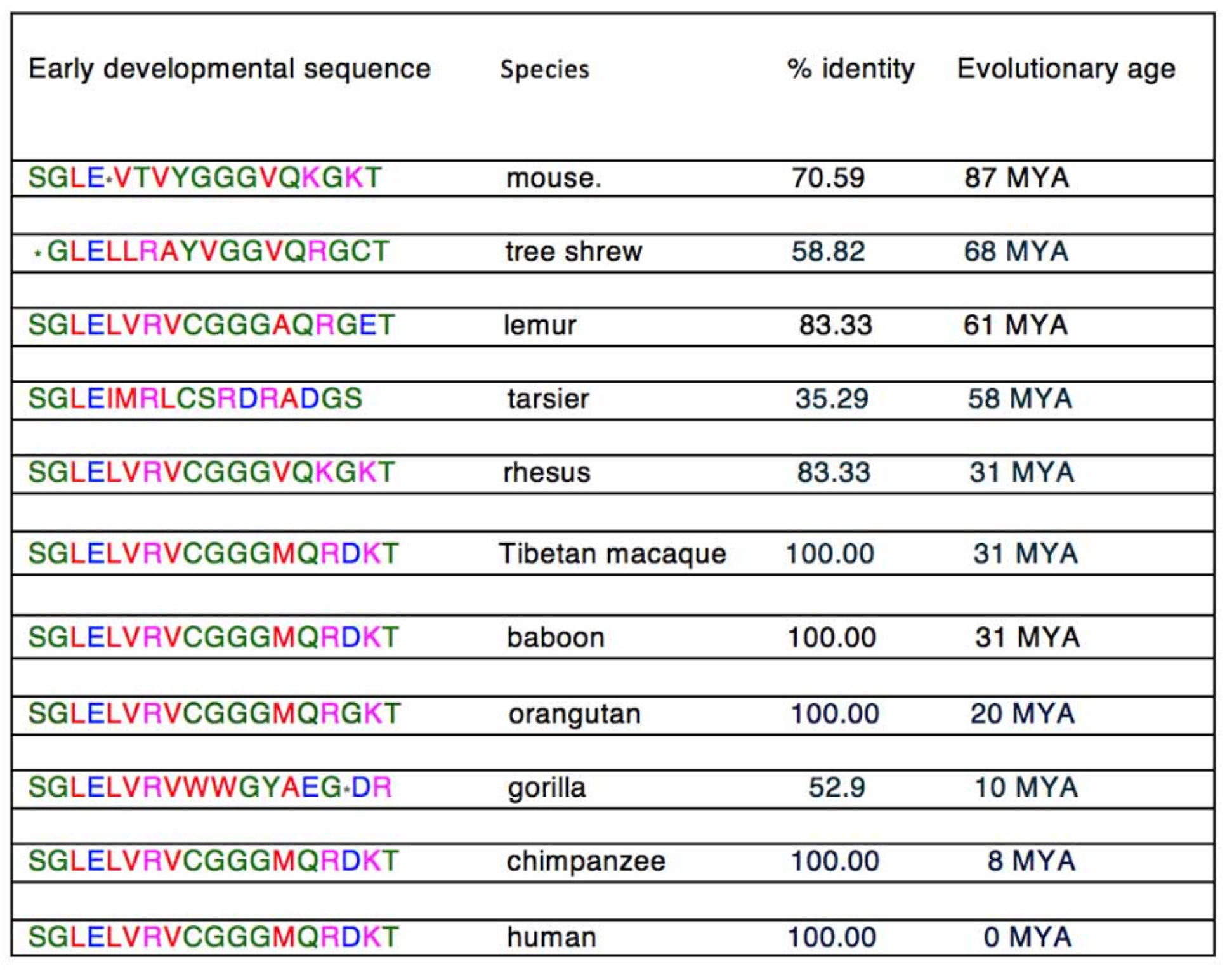
The evolutionary progression of the early developmental sequence in primates.

### Evolutionary development of the 107 aa ORF

The formation of the 107 aa ORF during primate evolution was analyzed. This readily provides a picture of how the 107 aa sequence grows during evolution. Specific primate species were chosen for analysis to demonstrate examples of random mutations occurring.

With the mouse sequence (Fig. 6), alignment of the translated 5’3’ Frame 3 aa sequence (shown in Fig. 4b) displays the SGLE**\***VTVYGGGVQKGKT sequence but no other significant contiguous stretches of aa residues with an identity to the human 107 aa. There are individual amino acid residues that show similarities, e.g., the mouse _2_CP, and _9_F, but they may have identity with the human107 aa by chance.

**Figure 6.**
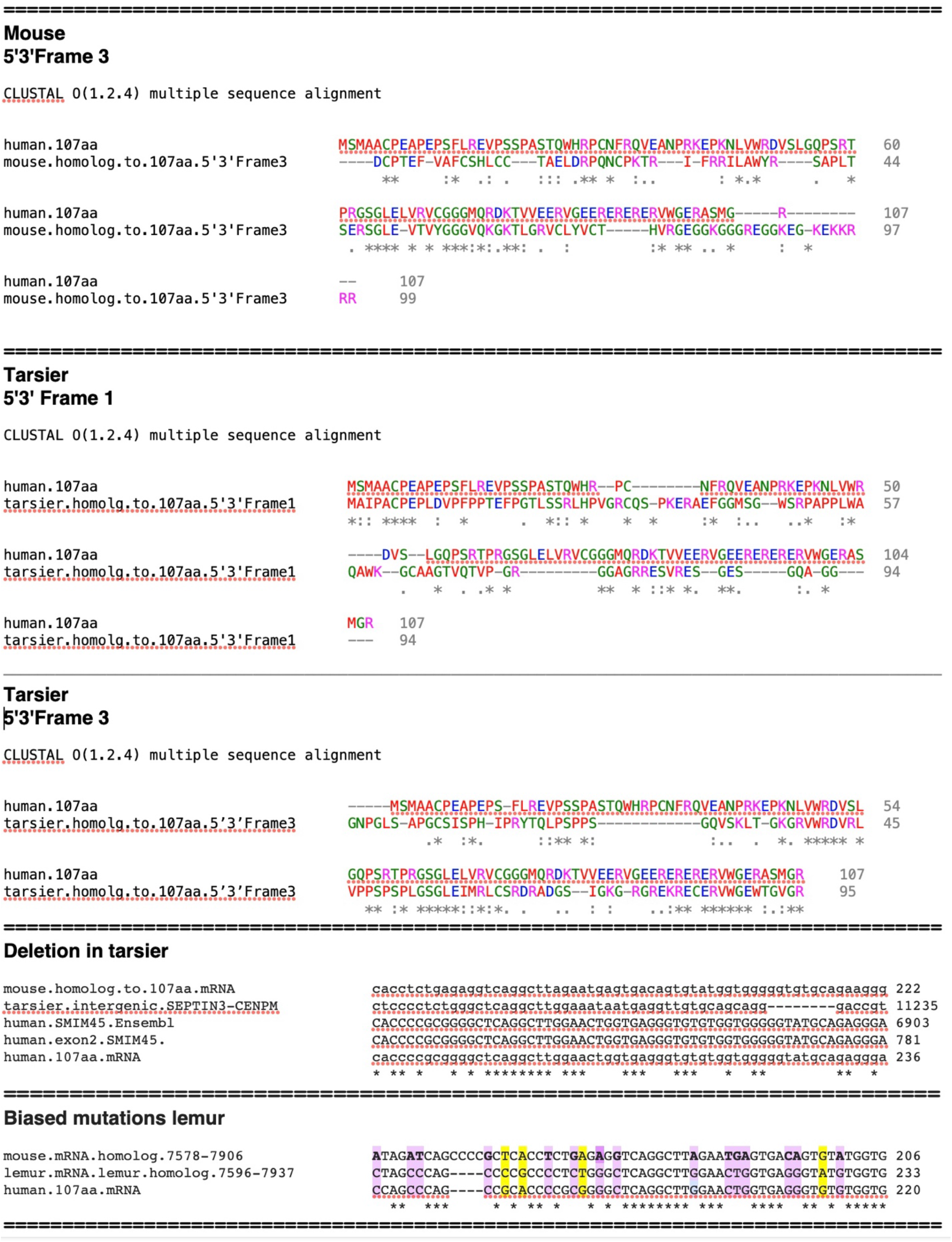
Development of the *SMIM45* 107aa ORF in the mouse and tarsier. The source of the mouse aa sequences is from reading frames of translated genomic nt sequences homologous to the 107aa mRNA sequence. The dash in the sequence _48_SGLE-VTVYGGGVQKGKT_64_ refers to a stop codon. The alignment of the human 107 aa sequence with the mouse 5’3’ Frame 3 aa (obtained from a translation of the mouse nt sequence homologous to the 107 aa mRNA), and, alignment of 5’3’ Frames 1 and 3 of the translated tarsier sequence. **Figure 6. bottom**,**” Deletion in tarsier 107 aa mRNA sequence”**. Nt sequence alignment show the deletion of nine bps in the tarsier nt 107 aa sequence. **Figure 6. Bottom**, **“Biased and random mutations”** shows mutations between the mouse, lemur and human sequences.

With further evolutionary development of the 107 aa with time, the Philippine tarsier (*Carlito syrichta*) sequence shows that it has progressed from that of the mouse and displays a methionine start aa residue with four contiguous aa residues of identity that are close to the start codon (Fig. 6, tarsier, 5’3’ Frame 1). There is also the beginning of development of aa identity in the middle of the sequence and at the C-terminal end (5’3’ Frame 3). The presence of segments of similarity in different regions of the tarsier sequence suggests that development of the 107 aa sequence progresses by formation of bps of identity in different genomic sections. Of interest, the human aa sequence, _73_GGGMQRDKT_81_, which is part of the early developmental sequence and present in the mouse genome, is absent in the tarsier homolog. This is due to point mutations and the deletion of the mouse nt sequence GGTGTGCA in the tarsier, but the sequence is present in the human 107 aa RNA (Fig. 6, bottom, “Deletion in tarsier”). This represents a random mutation specific to the tarsier and it is not found in progeny.

In analyzing nt sequences of genomic segments of the mouse and lemur [*Lemur catta* (Ring-tailed lemur)] that are homologous to that of the human mRNA sequence,15 mutations are evolutionarily fixed (highlighted in purple) in the lemur/human, compared to 4 random mutations (highlighted in yellow, Fig. 6, bottom “Biased and random mutations lemur”; these biased mutations may be selected and fixed by cellular regulatory mechanisms. The lemur predates the tarsier by approximately 7 MY, however, both are primitive primates and members of the prosimians. The rationale for choosing this particular genomic segment is that it shows a high similarity in nt sequence between species. Analysis in other regions of the lemur sequence where there is more sequence variability is difficult as some bp identities may be by chance. The alignment in the figure shows both fixed (biased) and random mutations that supports the concept of biased random mutations occurring during 107 aa ORF evolutionary development.

The rhesus sequence shows a significant increase in similarity at the amino terminal end compared to the tarsier. (Fig. 7, 5’3’ Frame1). However, similar to the tarsier, there is significant development of sequence identity in other sections; the 5’3’ Frame 2 shows larger stretches of contiguous sequence identity in the middle and carboxyl terminal regions. The elongation of tandem aa residues shows biased nearest neighbor additions.

**Fig. 7.**
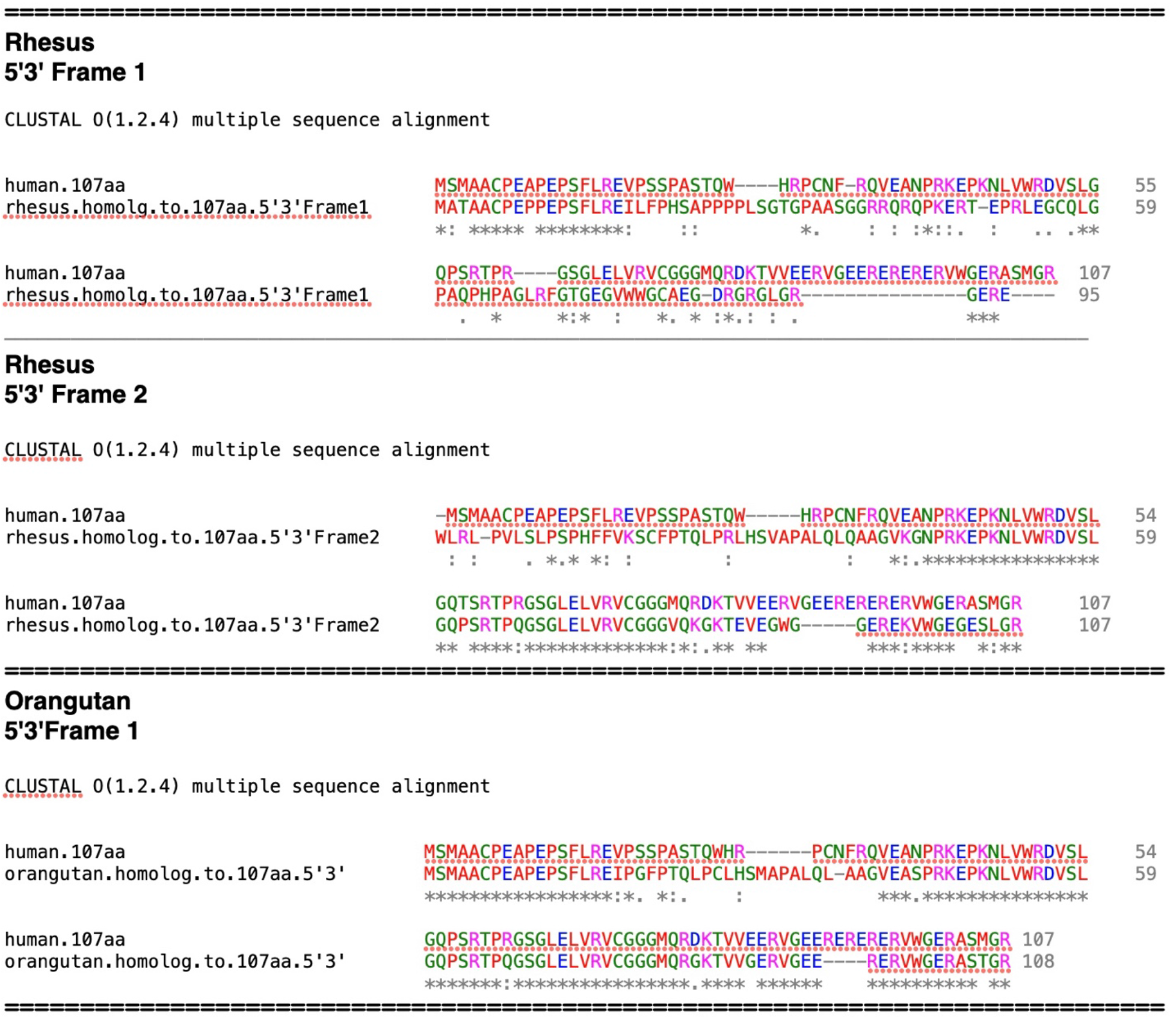
**Top.** Amino acid alignment of the rhesus sequence homologous to the 107 aa. The 5’3’ Frames 1 and 2 sequences are shown. **Bottom**, alignment the homologous 107 aa orangutan sequence with that of the human 107 aa.

The orangutan sequence displays a further increase in contiguous aa identity in all regions and does not display frame shift mutations (Fig. 7). The aa sequence identity compared to the human aa sequence is 79 %. However, a 13 bp insertion in the orangutan mRNA, which is not found in the rhesus, chimpanzee or human mRNAs, results in addition of the aa sequence **_30_**SMAPAL**_35_** (Fig. 7). This is due to a random mutation but, as with other random mutations mentioned, it is not carried over to progeny.

The chimpanzee sequence shows an almost complete human 107 aa ORF, except for the lack of ten C-terminal amino acid residues (VWGERASMGR**_107_**) that is present in the human sequence (Fig. 8, 5’3’ Frame 3). Of interest, this is due to a four bp tandem repeat AGAG, which is missing in the chimpanzee nt sequence, and results in a frame shift (Fig. 8, Bottom, “Alignment of chimpanzee nt sequence with human 107aa mRNA”, highlighted in turquoise). However, the chimpanzee 5’3’ Frame 2 aa alignment shows the presence of the ten C-terminal aa sequence in the chimpanzee genome. With the exception of three-point mutations, the absence of the small tandem repeat, AGAG distinguishes the chimpanzee 107 aa homolog from the human 107 aa. The tandem repeat may be due to slipped-strand mispairing (Levinson and Gutman 1987) that occurs at repeat sequences and constitutes a random process. Thus, the major difference between the chimpanzee and human aa sequences appears to be due to random mutations. Levinson and Gutman (1987) first proposed the concept of slipped-strand mispairing as a driving force in the evolution of DNA.

**Fig. 8.**
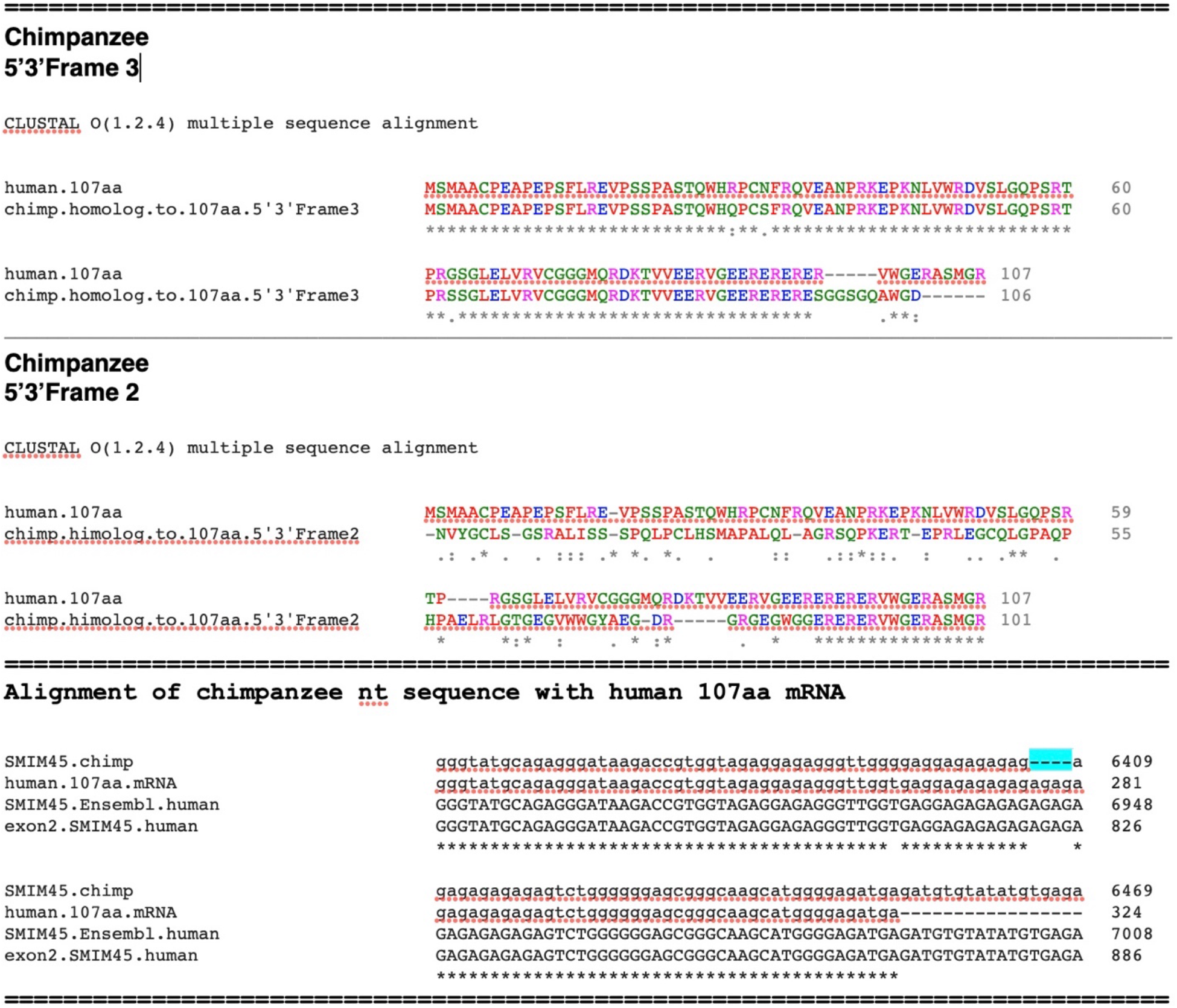
Alignment of the evolving 107 aa sequence in the chimpanzee with that of the human 107 aa. Displayed are 5’3’ Frames 2 and 3 of the chimpanzee homologous sequence. **Bottom**. An Alignment of segments of the chimpanzee *SMIM45* sequence, human *SMIM45* exon 2, the human *SMIM45* sequence from *Ensembl,* and the human 107 aa mRNA sequence. The alignment shows the absence of the 4 bp AGAG tandem repeat in the chimpanzee sequence (highlighted in turquoise).

Fig. 9 is a graphic representation that summarizes the evolutionary formation of the 107 aa ORF described in Figs 6-8. In Fig. 9, the open spaces between the NH_2_ and COOH ends represent random aa sequences, and the black dashed lines represent aa residues with identity in the 107 aa sequence. In the mouse*SMIM45* locus, shown is the the early developmental sequence flanked by random sequences. In the tarsier, the 107 aa sequence development shows a methionine start and formation aa sequence identity at the NH2 and COOH terminal ends. Growth of the 107 aa sequence continues with nearest neighbor additions at the three separate regions, the NH2 end, the middle and the COOH terminal end in the rhesus and orangutan loci. The chimpanzee shows the joining of segments to provide a contiguous 107 aa sequence with the exception of the COOH terminal end that is missing 10 aa residues of the human107 aa sequence due to a frame shift. This is corrected in the human 107 aa sequence with addition of a tandem four base pair repeat.

**Fig. 9.**
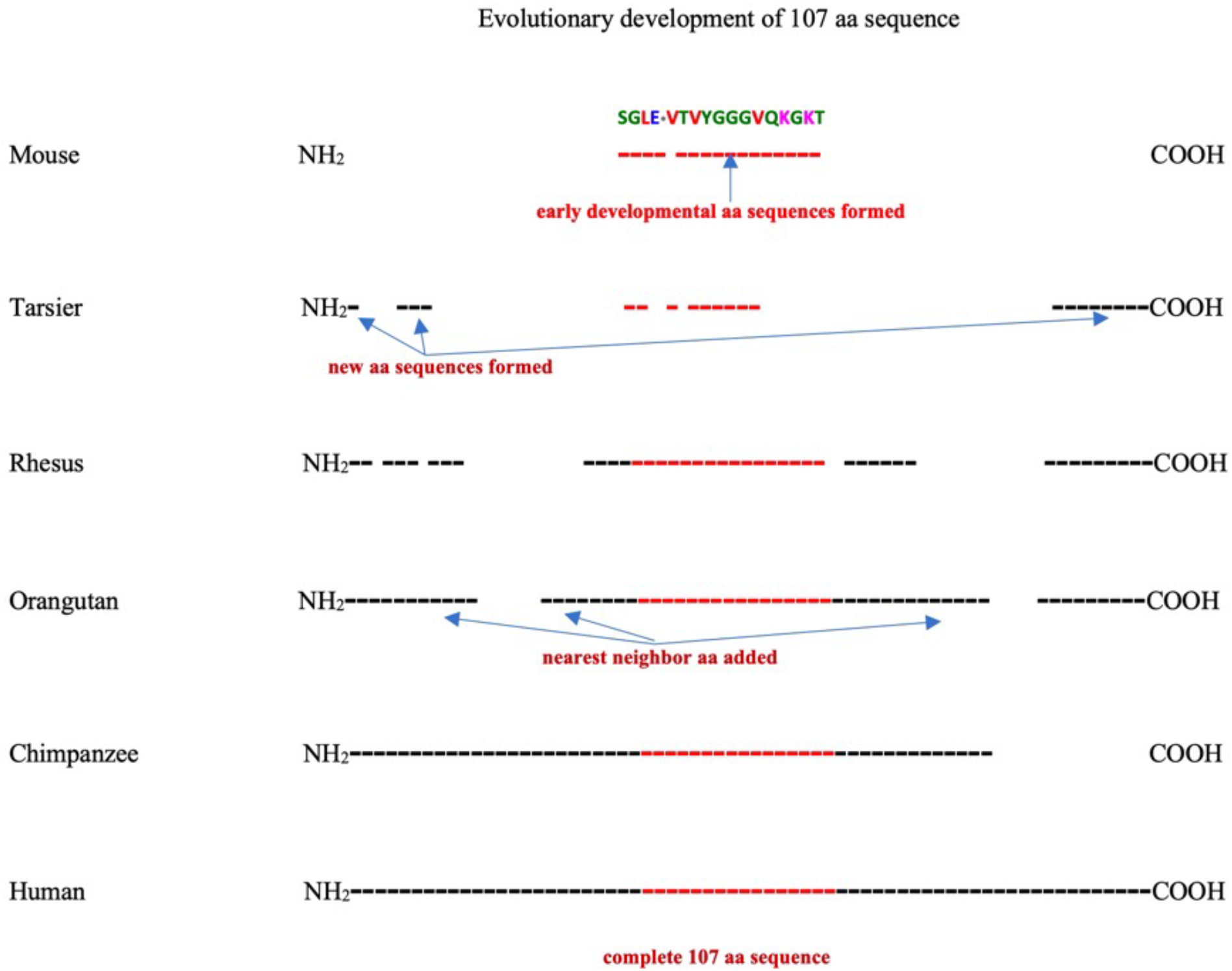
Diagrammatic representation of the evolutionary formation of the human 107 aa sequence. The open spaces represent regions with no aa sequence identity to the 107 aa human 107 aa and are considered random sequences (note, there are individual aa residues with similarities to the human 107 aa sequence in these regions, but we can not determine if the identity is random or not). The dashed lines represent aa that have identity to the human 107 aa sequence. The representation depicts random mutations in the chimpanzee (COOH end) and in the tarsier homolog to the mouse SGLE**\***VTVYGGGVQKGKT sequence.

### Search for an evolutionary root organism

<u>T</u>he mouse *SMIM45* 107 aa early developmental sequence has been used as a basis for the evolutionary formation of the 107aa sequence as there is strong statistical evidence that it represents part of the 107 aa. There is weaker evidence for the presence of the sequence in several ancestors of the mouse. Nevertheless, certain <u>a</u>ncestors close to the evolutionary age of the mouse were analyzed in an effort to determine species root origins of the 107 aa sequence. Species investigated are shown below in evolutionary chronological time (Miller et al. 2007; Phillips and Fruciano 2018: Warren et al. 2008):

Mouse (87 MYA), rabbit, cat, cow, mole, elephant (103 MYA), koala (164 MYA) and platypus (166 MYA)

The analysis was focused on the early developmental sequence, both its nt and aa sequences as well as a display of synteny with the 68 aa genomic location.. This provided a means to follow the evolutionary progression of the 107aa in species ancestral to the mouse; analysis by using only the entire 107 aa and nt sequences was complicated by random sequences.

The *SMIM45* gene sequences of the rabbit, cat, cow, mole and elephant species show partial aa identity with the early developmental human sequence SGLELVRVCGGGMQRDK. (Supplementary file 1, Figs. S2-S6). The elephant *SMIM45* gene sequence is of significant interest as the identity of the early developmental aa sequence with that of the human sequence is 44%, and the nt sequence that translates to the early developmental aa sequence is 79% (Supplementary file 1, Fig. S6). This demonstrates that root formation of the human SGLELVRVCGGGMQRDK sequence dates to an ancestral species whose evolutionary appearance precedes 103 MY (the evolutionary age of the elephant). The ancestral species of the elephant, the koala (Phillips and Fruciano 2018) and platypus (Nilsson et al. 2010) display no significant identity to either the aa or nt sequence of the early developmental motif (Supplemental file S1, Figs. S7 and S8). This suggests the root species of origin of the 107 aa ORF is of an evolutionary age between that of the koala and elephant. Finding a species that shows a marginal “twilight zone” identity with the early development sequence would likely represent a near root organism, but in a search of species genomes we have not been able to find one. The opossum (*Monodelphis domestica*, gray short-tailed opossum) is a species that is evolutionarily between the elephant and the koala (Nilsson et al, 2010); however, the *CENPM* or *SMIM45* genes have not been annotated in the opossum as sequences in the *CENPM*-*SEPTIN3* region have not been established; this precludes an analysis of the 107 aa early development sequence in this species. Another species whose evolutionary origin is between the elephant and koala is the *Elephantulus edwardii* (Cape elephant shrew) but similar to the opossum, the Cape elephant shrew *SMIM45* gene sequence has not been determined.

In Supplementary file 1, Fig. S9 is a color coordinated DNA sequence alignment that shows the totally conserved bases in the early developmental nt sequence during evolution of the species analyzed. The alignment demonstrates a large bias towards guanines that remain totally constant during the evolution of the species, for example, 9 out of 21 Gs remain totally conserved compared to 1 out of 7 As. This poses the interesting question of why guanines are far less likely to mutate (be subject to errors in DNA replication) than other bases during the evolution of the early developmental sequence.

## Discussion

By comparing the progression of two *SMIM45* ORFs during evolutionary time a sharp contrast in sequence conservation vs sequence change is found side by side. The 68 aa ORF, a microprotein, exhibits strict regulation of bp conservation, e.g., the ORF shows 18 bps that differ between the mouse and human sequences, but there is only one bp substitution that results in a change in an aa residue; the other 17 mutations involve the wobble position. This implies an ancient regulatory element that protects against mutations that alter the ORF. Why the 68 aa open ORF is 100% conserved over ∼75 MY during primate evolution offers an interesting question. It has a predicted alpha transmembrane domain (Supplemental file S1, Fig. S1) (Hallgren J. et al. 2022) and its DNA is GC rich, but a protein product has not been isolated and characterized. On the other hand, one of the two genes that shows synteny with *SMIM45*, *SEPTIN3* (Fig. 1) is also ultra-conserved but has a known function, i.e., that of binding proteins associated with autophagy (Tóth et al. 2022).

A model of the evolutionary development of the 107 aa sequence can be visualized with an initial selection of a bp in a genomic region that displays a random sequence. This constitutes a putative starter seed bp that is evolutionarily fixed. The next stage consists of nearest neighbor sequence growth, where in the example of the mouse forms the bp sequence encoding the aa sequence SGLE**\***VTVYGGGVQKGKT. Other putative root bps in different regions of the *SMIM45* gene also expand with nearest neighbor additions. Growth is not by continuous elongation from one region, e.g., the N-terminal. The full human 107 aa sequence forms when all nearest neighbor positions are joined. There is a bias in seed base pair selection and in nearest neighbor sequence growth. Random mutations also occur during the process of aa chain growth that result in point mutations and frame shifts during 107 aa development, however these are corrected as they are not found in descendant species. Regulatory elements dictate the initial seed bps, the specific nearest neighbor formation, and the protection of the growing sequence from disruptive mutations.

A study of the partial evolutionary progression of a number of protein genes in primates was described (Broeils et al. 2023). One example is the *MYEOV* (myeloma overexpressed) gene that shows a frame shift and one base pair change in the first codon creating a Met initiation at the N-terminal end that distinguish the chimpanzee sequence from that of the human sequence. Data were not presented for the determination of a possible early developmental stage of this gene. However, the *MYEOV* mRNAs from various primate species show a very significant development of the *MYEOV* mRNA sequence in the rhesus macaque, suggesting a rapid nt sequence development from the New World primates or the primitive lower prosimian primates.

Biased random mutations can be considered analogous to a biased random walk, a concept used to describe the migration of bacterial cells, leucocytes and other cells in space during chemotaxis (Macnab and Koshland Jr 1972; Koshland DE Jr 1980; Sourjik et al 2012). A diagram of bacterial chemotaxis is shown in Supplementary file S1, Fig. S10. In chemotaxis, the bias is movement towards a “goal”, e.g., a high concentration of nutrient, but there are also random movements away from the nutrient. Biased mutations during evolution as presented here lead to the formation of the human 107aa open reading frame, the “goal”, but with random mutations that represent drift; these can be considered analogous to small movements in random directions in chemotaxis. In both cases, the randomness is corrected. The progression with time of ORF development is the variable analogous to movement in space in chemotaxis. If enough random mutations can be pinpointed for statistical significance, a mathematical model of a biased mutational process, analogous to a mathematical model of a biased random walk, may be helpful in better understanding the mutational process associated with open reading frame development during evolution. Mathematical models for chemotaxis have been presented before to better access biased random walks (Codling et al. 2008) (Liu et al. 2023).

## Conclusions

Evolutionary analysis of the human 107aa ORF indicates that it was predetermined over 100 MYA. Formation begins in non-primate species and initially detected is a short early developmental sequence. In primates, sequence formation progresses in different regions in the *SMIM45* locus; thus, growth is not by continuous elongation from one region, e.g., the N-terminal. This pattern of development may also pertain to evolution of human *de novo* protein genes. The function of the 107 aa ORF is uncertain. An et al (2023) proposed a role in human fetal brain development, however further experimental work with protein isolations is needed. It is possible that the 107 aa sequence is expressed transiently and only in certain tissues. Clarification is needed to understand why the SMIM45 transcript variant 1 mRNA (NM_001395940.1), which is transcribed and can code for both the 68 aa and the 107 aa nt sequences, shows no translation of the107 aa nt sequence. This may change, but as it stands, the current lack of evidence for a protein product, balanced against the stringently regulated developmental process offers a paradox. The RNA transcript NM_001395940.1 appears to fall into a special category that can be described as producing a translated microprotein (68 aa) and as a “transcript of unknown function” (the 107 aa ORF).

## Supporting information

Supplement S1 copy

## Acknowledgements

I thank Dr. Alan Tucker, Department of Applied Mathematics and Statistics, Stony Brook University, New York for statistical analyses, Dr Christopher Gilbert, Department of Anthropology, Hunter College, City University of New York for recent data on primate evolutionary ages (molecular divergence), and Dr. Matthew Philips, School of Biology and Environmental Science, Brisbane, Australia for the evolutionary aspects of the marsupials.

