## Supplement S1 copy for "Evolutionary formation of a human *de novo* open reading frame from a non-primate non-coding genomic region that proceeds via biased random mutations"

Nicholas Delihas

https://orcid.org/0000-0002-1704-2587

Department of Microbiology and Immunology
Renaissance School of Medicine
Stony Brook University
Stony Brook, NY 11794 USA


Tel. # 001 631 286-9427

#### Supplementary file S1

**Fig. S1.** Predicted transmembrane domain of the 68 aa sequence. Data obtained using the DeepTMHMM prediction program of Hallgren et al. (Hallgren J. et al. bioRxiv preprint doi: https://doi.org/10.1101/2022.04.08.487609).

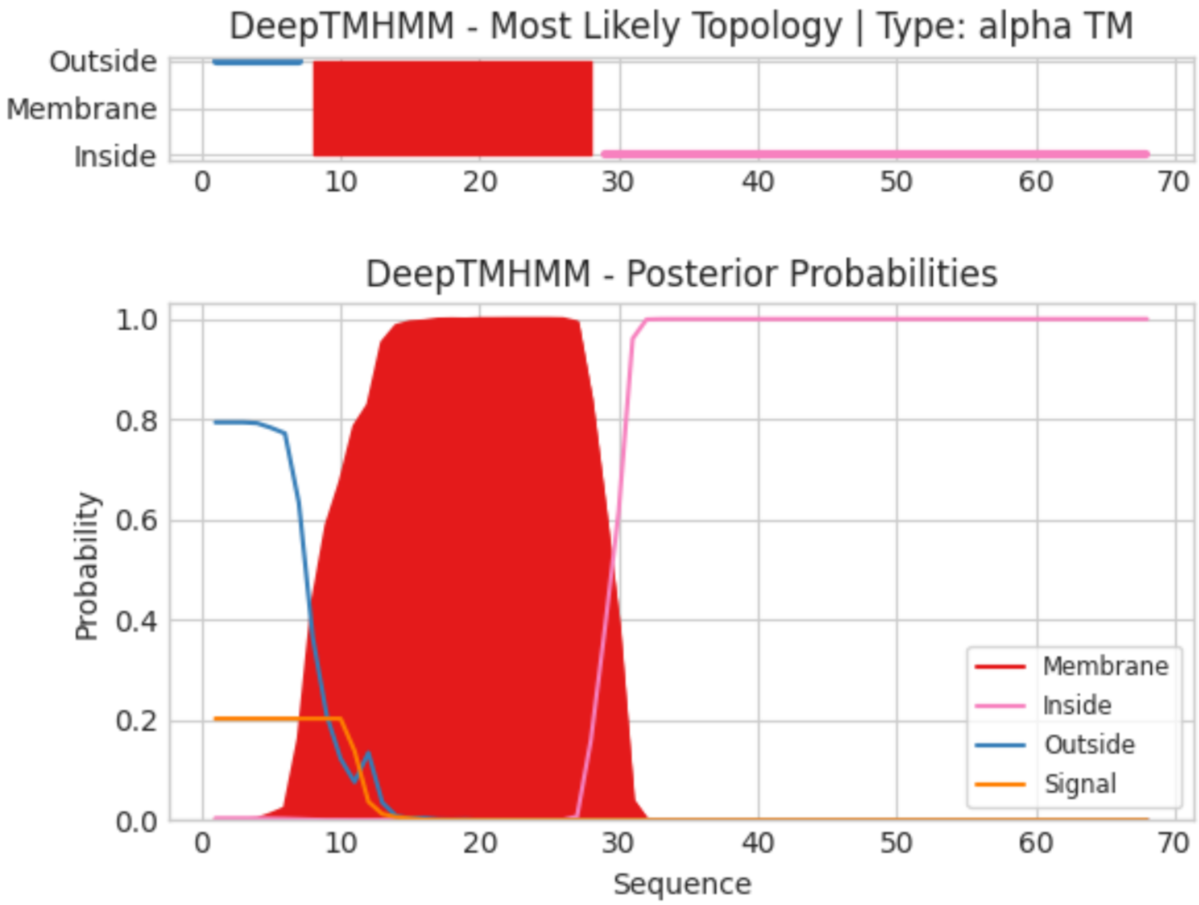

==================================================================

### Rabbit

**Fig. S2**. Alignment of the rabbit early developmental aa sequence with the human sequence. **Top.** The rabbit aa sequence shows 56% identity and **Bottom.** 83% nt sequence identity. Point mutations and a frame-shift mutation results in a lower aa identity to the human sequence relative to that of the mouse (67% identity).

##### Top

```
#  Percent Identity Matrix - created by Clustal2.1

     1: rabbit      56.25
     2: human      100.00

CLUSTAL O(1.2.4) multiple sequence alignment

rabbit       SGLEVGRVWGGGLRVR- 16
human        SGLELVRVCGGGMQRDK 17
             ****  *  ***::
```

##### Bottom

```
# Identity:      45/54 (83.3%)
# Similarity:    45/54 (83.3%)
# Gaps:           1/54 ( 1.9%)
# Score: 228.0
#
#
#=======================================

Rabbit.homolo       1 tcaggcttggaagtggggagggtttggggtgggggcctgtagaggg-taa      49
                        ||||||||||||.|||.||||||.||.||||||||..||.|||||| |||
SGLELVRVCGGGM       1 tcaggcttggaactggtgagggtgtgtggtgggggtatgcagagggataa      50

Rabbit.homolo      50 gagc      53
                        ||.|
SGLELVRVCGGGM      51 gacc      54
```

==============================================================

### CAT

**Fig. S3. Top**. Identity and alignment of the 5’3’Frame1 aa translation of the cat homolog to the early development aa sequence showing a 29% identity. **Bottom**. Nt sequence alignment of the cat homolog to the early development with 107 aa mRNA early development nt seq. Point mutations and a frame shift mutation in the cat homolog to the SGLELVRVCGGGMQRDKT nt sequence deceases the similarity in aa sequence.

#### Top

```
#  Percent Identity Matrix - created by Clustal2.1

     1: 5'3'Frame1.cat              29.41
     2: SGLELVRVCGGGMQRDKT.human   100.00

CLUSTAL O(1.2.4) multiple sequence alignment

5'3'Frame1.cat                SDLAMVSVCDRVCRGVR-      17
SGLELVRVCGGGMQRDKT.human      SGLELVRVCGGGMQRDKT      18
                              *.* :* **.   :   :
```

#### Bottom

```
# Identity:      43/54 (79.6%)
# Similarity:    43/54 (79.6%)
# Gaps:           1/54 ( 1.9%)
# Score: 234.0
#
#
#=======================================

cat.homolog.t      1 tcagacttggcaatggtgagcgtgtgtgac-agggtgtgcagaggggtga      49
                       ||||.|||||.|.|||||||.|||||||.. .||||.|||||||||.|.|
human.SGLELVR      1 tcaggcttggaactggtgagggtgtgtggtggggtatgcagagggataa      50

cat.homolog.t     50 gacc      53
                     ||||
human.SGLELVR     51 gacc      54
```

==================================================================

### Cow

**Fig. S4. Top.** Alignment of the cow early developmental aa sequence with the human sequence shows a low aa identity, 23% **Bottom.** The cow nt sequence a large number of point mutations that result in poor identity to the human sequence, 52%.

#### Top

```
#  Percent Identity Matrix - created by Clustal2.1

     1: human       100.00
     2: mouse        70.59
     3: cattle.      23.08

CLUSTAL O(1.2.4) multiple sequence alignment


human                   SGLELVRVCGGGMQRDKT-----   18
mouse                   SGLE-VTVYGGGVQKGKT-----   17
cattle.5’3’Frame1       ---S-LGRLG-GLSSEGTERLGS   18
                           . :   * *:.   *
```

#### Bottom

```
#  Percent Identity Matrix - created by Clustal2.1

     1: cattle.homolog.to.nt.of.SGLELVRVCGGGMQRDKT  51.85
     2: human.seq.tansl.to.SGLELVRVCGGGMQRDKT      100.00
     3: mouse.seq.transl.to.SGLE*VTVYGGGVQKGKT      79.63


CLUSTAL O(1.2.4) multiple sequence alignment


cattle  tcataactaggacggttgggaggcttgtcgtctgagggtacggagagattggggtcc 57
human   tcaggcttggaactggt-gagggtgtgtggtgggggtatgcagagggataagacc-- 54
mouse   tcaggcttagaatgagt-gacagtgtatggtgggggtgtgcagaagggtaagact-- 54
        ***    * * *    * *   *  * * **  * *  * * **  * *  *
```

==================================================================

### Mole

**Fig. S5.** Alignment early developmental sequence of the mole with that of the human sequence. **Top.** The aa alignment shows a 38% identity. **Bottom.** The mole nt sequence shows point mutations, a frame-shift mutation and the absence of the first six bp at the 5’; end.

#### Top

```
#  Percent Identity Matrix - created by Clustal2.1

     1: SGLELVRVCGGGMQRDKT.nt.seq 100.00
     2: 5'3'Frame1                37.50

CLUSTAL O(1.2.4) multiple sequence alignment


SGLELVRVCGGGMQRDKT.nt.seq      SGLELVRVCGGGMQRDKT     18
5'3'Frame1                     LGLECVSVWGEQWAQT--     16
                                *** * * *    :
```

#### Bottom

```
# Length: 56
# Identity:      34/56 (60.7%)
# Similarity:    34/56 (60.7%)
# Gaps:          10/56 (17.9%)
# Score: 91.5
#
#
#=======================================

SGLELVRVCGGGM      1 tcaggcttggaactggtgagggtgtgtg-gtg-ggggtatgcagagggat     48
                           .|||..|||  |||.|||||.| ||| |||||..||||.|||..
mole.homolog.      1 ------ctgggcctg--gagtgtgtgagtgtgtggggtgagcagtgggcc     42

SGLELVRVCGGGM     49 aagacc     54
                     .||||.
mole.homolog.     43 cagaca     48
```

===================================================================

### Elephant

**Fig. S6.** Alignment of the early developmental sequence of the elephant with that of the human sequence. **Top.** Early developmental aa sequence shows 44% identity. **Bottom**. The elephant nt sequence homologous to the early developmental sequence aligned with the human early developmental sequence shows point mutations and a frame-shift mutation. The nt sequence identity is comparable to that of the mouse, 78% elephant, 81% mouse.

##### Top

```
#   Percent Identity Matrix - created by Clustal2.1

     1: 5'3'Frame1.elephant 100.00
     2: early.dev.seq        44.44


CLUSTAL O(1.2.4) multiple sequence alignment


5'3'Frame1.elephant      SGLEWVRWVWLGVCRRAS 18
early.dev.seq            SGLELVRVCGGGMQRDKT 18
                         **** **     *: *   :
```

##### Bottom

```
# Length: 56
# Identity:     44/56 (78.6%)
# Similarity:   44/56 (78.6%)
# Gaps:          2/56 ( 3.6%)
# Score: 160.0

#=======================================

elephant.exte      1 tcaggtttagaatgggtgaggtgggtgtggctgggggtgtgcagaagggc     50
                     |||||.||.|||..||||||| |.|||||| |||||||.||||||.||..
SGLELVRVCGGGM      1 tcaggcttggaactggtgagg-gtgtgtgg-tgggggtatgcagagggat     48

elephant.exte     51 aagccc     56
                     |||.||
SGLELVRVCGGGM     49 aagacc     54
```

==========================================================

### Koala

**Fig. S7.** Alignment of the koala *SMIM45* gene sequence with the 68 aa and 107 aa mRNAs, and the RNA that codes for SGLELVRVCGGGMQRDKT.

The alignment shows the 68 aa and 107 aa co-align with the koala SMIM45 gene indicating that there is no display of synteny and the alignment appears random.

```
68aa          ---------a tgccgcactt cctggactgg ttcgtgccgg tctacttggt
koala gene    gccagggcca tgccgcattt tctggactgg tttgtgcccg tctatctcat
107aa         gcca--gccc agccgcaccc cgcggg---- ---------- ----------
SGLELVRVCG    ---------- ---------- ---------- ---------- ----------

              catctcggtc ctcattctgg tgggcttcgg cgcctgcatc tactacttcg
              gatctccatc ctcatcctgg ttggtttcgg ggcctgtatc tactacttcg
              ---------- ---------- ---------- ---------- ----gctcag
            | ---------- ---------- ---------- ---------- ------tcag

              agccgggcct gcaggaggcg cacaagtggc gcatgcagcg ccccctggtg
              agcccggttt gcgggaagcc cacaagtggc gaacacagag acccctgctg
              gcttggaact ggtgagggtg tgtggtgggg gtatgcagag g---------
              gcttggaact ggtgagggtg tgtggtgggg gtatgcagag g---------

              gaccgcgacc tccgcaagac gctaatggtg cgcgacaacc tggccttcgg
              gaccggaact ttcgcaagac cctgatgatc caggacaacc tggcatttgg
              ---------- --gataagac cgtggtagag ga-------- ----------
              ---------- --gataagac c--------- ---------- ----------

              cggcccggag gtctg----- ---------- ---------- ----------
              gagccccgat gtctgatggg tctggggccg agacagatgg ggggacaggg
              -------gag ggttggtg-- ---------- ---------- ----------
              ---------- ---------- ---------- ---------- ----------
```

==================================================================

### Platypus

**Fig. S8.** Alignment of a section of the platypus genomic sequence between genes *CENPM* and *SEPTIN3* with the human 107 aa mRNA, 68 aa mRNA, the RNA that codes for SGLELVRVCGGGMQRDKT and the uncharacterized platypus locus LOC114816411.

*SMIM45* has not annotated in the platypus genome. However, the 68 aa mRNA sequence aligns within the *CENPM -SEPTIN3* region at position 11188bp, and also at position 2469 of LOC114816411 (highlighted in turquoise). The 107 aa mRNA aligns primarily at the region near 1320 bp of the *CENPM -SEPTIN3* region, which is a very different genomic location: in addition, sections of the107 aa sequence are scattered in different locations indicating random identities. This thus shows a random sequence alignment of the 107 aa RNA and the early developmental sequence as well as the absence of synteny between the 107 aa and 68 aa mRNA sequences. The alignment also shows that LOC11481641 contains the 68 aa sequence, and suggests that the *SMIM45* gene is present in platypus and corresponds to LOC11481641.

**68 aa nt sequence alignment with platypus genome**

```
human.107aa.mRNA                            ------------------------------------------------------------    324
SGLELVRVCGGGMQRDKT.nt.seq                   ------------------------------------------------------------    54
CENPM.start.Septin3.start.47467892-47491106 ccctcacgctgtgcagcggcggccaccatgccccacttcctggactggttcgtgcccgtc    11220 11188 bp
platypus."SMIM45"Loc114816411               ccctcacgctgtgcagcggcggccaccatgccccacttcctggactggttcgtgcccgtc    2507. 2469 bp
human.68aa.mRNA                             ---------------------------ATGCCGCACTTCCTGGACTGGTTCGTGCCGGTC    33
```

//////////////////////////////////////////////////////////////////////////////////////////////////////////////////

**107 aa nt sequence alignment with platypus genome**

```
human.107aa.mRNA                            cgaagcca-----------------------acccaaggaaagaacctaagaacctcg-t    143
SGLELVRVCGGGMQRDKT.nt.seq                   ------------------------------------------------------------    0
CENPM.start.Septin3.start.47467892-47491106 cgcccacagtcacacagccgacaagtggcagagccgggaggcgaactcaggagccctgac    1320
platypus."SMIM45"Loc114816411               ------------------------------------------------------------    0
human.68aa mRNA                             ------------------------------------------------------------    0


human.107aa.mRNA                            ttggagggatgtcagcttgggccagc--ccagccgcaccccgcggggctcaggcttgg--    199
SGLELVRVCGGGMQRDKT.nt.seq                   ------------------------------------------------tcaggcttgg--    10
CENPM.start.Septin3.start.47467892-47491106 tccaaagcccgtgctctttccactgcgccccgctgcttctctatatgctctctatggagg    1380
platypus."SMIM45"Loc114816411               ------------------------------------------------------------    0
human.68aa                                  ------------------------------------------------------------    0
```

**Fig.S9.** Displayed is the alignment of the early developmental sequences from genomes of various species showing color coordinated DNA bases. This diagram readily shows that guanine bases of the early developmental sequence are far more conserved relative to the other bases during evolution, for example, there are 9 out of 21 Gs that remain totally conserved compared to 1 out of 7 As. 2 Ts out of 9, 0 out of 2 As.

The SGLELVRVCGGGMQRDKT nt sequence is G rich:
A(20%) T(23%) G(46%) C(11%)

Data was. obtained using  MAFFT online service: multiple sequence alignment, interactive sequence choice and visualization,, http://msa.biojs.net/ **(**Katoh, Rozewicki, Yamada (2019) MAFFT online service: multiple sequence alignment, interactive sequence choice and visualization. *Briefings in Bioinformatics* **20**:1160-1166; Kuraku, Zmasek, Nishimura, Katoh 2013 (*Nucleic Acids Research* **41**:W22-W28) aLeaves facilitates on-demand exploration of metazoan gene family trees on MAFFT sequence alignment server with enhanced interactivity)

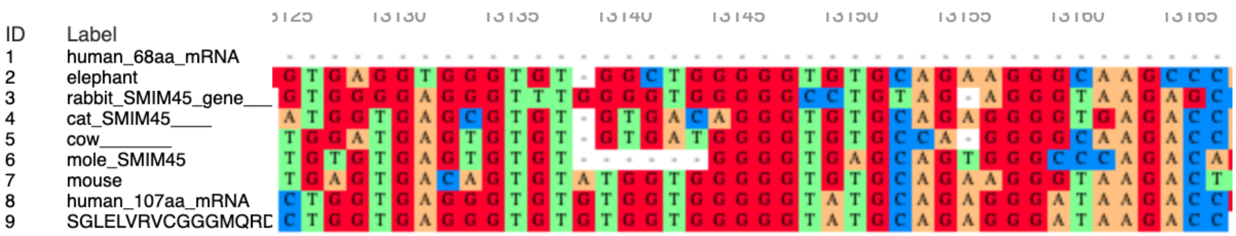

**Fig. S10.** Diagram of a cell moving towards an attractant with random movements in different directions.

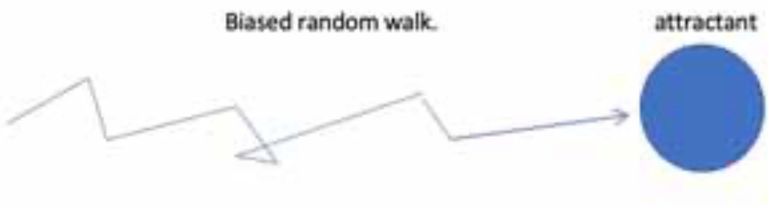


CLUSTAL O(1.2.4) multiple sequence alignment

cattle
